# Aggression is induced by resource limitation in the monarch caterpillar

**DOI:** 10.1101/2020.08.11.247346

**Authors:** Joseph Collie, Odelvys Granela, Elizabeth Brown, Alex C. Keene

## Abstract

Food represents a limiting resource for the growth and developmental progression of many animal species. As a consequence, competition over food, space, or other resources can trigger territoriality and aggressive behavior. Throughout their early development stages, insect larvae eat voraciously and limited food availability can potently impact their viability through metamorphosis. In the monarch butterfly, *Danaus plexippus,* caterpillars feed predominantly on milkweed, raising the possibility that access to milkweed is critical for growth and survival. Here, we characterize the role of food availability on aggression in monarch caterpillars. We find that monarch caterpillars display stereotyped aggressive lunges that increase during development, peaking during the 4^th^ and 5^th^ instar stages. Detailed behavioral analysis reveals that aggressive actions are most likely to occur when the target is feeding and increases the probability that the target will leave the food source. To determine the relationship between food availability and the initiation of an aggressive encounter, we provided groups of caterpillars differing amounts of food availability and measured aggressive behavior. The number of lunges toward a conspecific caterpillar was significantly increased under conditions of low food availability, suggesting resource defense may trigger aggression. We find that aggression occurs independently of light, suggesting the visual system is dispensable for the induction of aggression. These findings establish monarch caterpillars as a model for investigating interactions between resource availability and aggressive behavior under ecologically relevant conditions and set the stage for future investigations into the neuroethology of aggression in this system.

## Introduction

Resource competition is a primary driver of behavioral evolution that shapes interactions within conspecifics and between species (Peiman and Robinson, 2010; Brumm and Ritschard, 2011). Animals often compete for diverse resources throughout their lifetimes including territory, mates, and food acquisition. Aggression and territoriality are present from invertebrates through primates, and many of the molecular and neural mechanisms governing aggression are conserved across phyla (Thomas, Davis and Dierick, 2015). Food availability represents a critical resource that is particularly important in early life, as insufficient food consumption can potently affect developmental progression and brain development (Ormerod *et al.*, 2017; Keene *et al.*, 2019). Further, even short periods of food restriction can induce mortality during early life when animals often have lower energy stores or are undergoing developmentally specified rapid growth (Dawirs, 1984; Tu and Tatar, 2003). Therefore, while resource competition and aggression are often studied in adult animals, these behaviors are also likely to be adaptive early in life.

Aggressive behavior represents a heterogeneous behavior that consists of offensive aggression, often to acquire resources, and defensive aggression, to protect against predation (Takahashi and Miczek, 2013; Ormerod *et al.*, 2017; Wranghama, 2017). Aggression has been widely documented in eusocial and solitary insects where its function is used to defend territories, establish social hierarchies, and compete for food. For example, multiple species of butterflies have been studied for their territoriality, where males defend sunspots that attract potential mates (Davies, 1977; Wickman and Wiklund, 1983; Stutt and Willmer, 1998; Bergman, Olofsson and Wiklund, 2010). Under laboratory conditions, the fruit fly, *Drosophila melanogaster*, has become a leading model for studying the molecular mechanisms underlying aggression, which can be experimentally induced in the laboratory by pairing male flies in an arena with food (Chen *et al.*, 2002).

The standardization of this system, combined with powerful genetics has allowed for dissection of genes and neural circuits regulating aggressive behavior (Kravitz and Fernandez, 2015; Anderson, 2016; Hoopfer, 2016). Together, intraspecies aggression can be studied in diverse insect species under natural and laboratory conditions, providing the opportunity to investigate how environmental and social factors contribute to this behavior.

Food availability in early life may be particularly important in insect larvae that consume food voraciously throughout most larval stages (Edgar, 2006; Tennessen and Thummel, 2011). Nutritional restriction during larval stages has been shown to delay larval development as well as reduce adult body size, reproductive performance, and lifespan (BEADLE, TATUM and CLANCY, 1938; Dmitriew and Rowe, 2011; Runagall-Mcnaull, Bonduriansky and Crean, 2015; Poças, Crosbie and Mirth, 2020). Food consumption in larvae is essential to accommodate the growth necessary to achieve metamorphosis. Larvae from many species are social, and often compete for resources (Martin L. Cody, Robert H. MacArthur, 1975; Kaplan and Denno, 2007) For example, acoustic territoriality in Lepidoptera caterpillars (Yack, Smith and Weatherhead, 2001) and cannibalism in *Drosophila* larvae, beetles, and other insects (Dickinson, 1992; Vijendravarma, Narasimha and Kawecki, 2013; Orrock, Connolly and Kitchen, 2017; Huang, Miller and Walker, 2018) suggest the presence of aggression in early developmental stages. The social interactions and competition for resources raise the possibility that limited food availability triggers aggression during the early stages of insect development.

The monarch butterfly, *Danaus plexippus,* is a model for investigating the mechanisms underlying migration, navigation, and circadian function (Reppert, Gegear and Merlin, 2010; Zhan *et al.*, 2011; Tomioka and Matsumoto, 2015). Monarch caterpillars predominantly feed on milkweed and often strip entire plants bare of leaves over a two-week period. In many locations, milkweed is only available for part of the year, placing a significant constraint on monarch development (Yang and Cenzer, 2019). These ethological constraints suggest behavioral repertoires may be present in monarch caterpillars to acquire and defend resources.

To examine whether caterpillars display aggressive behavior, we established a group aggression assay and quantified the presence of aggressive lunges under a number of conditions, as well as the effect of attacks on target conspecifics. Here, we describe aggressive behavior in monarch caterpillars, which occurs during later stages of caterpillar development when resource competition may increase in the wild. Further, limiting food availability promotes aggression, suggesting that this represents a form of resource defense. These findings suggest monarch caterpillars may be used as a model to investigate how resource limitation contributes to regulation of aggressive behavior, and subsequent effects on the animals’ brain development and function.

## Methods

### Animal acquisition and husbandry

Monarch caterpillars were field-collected between May and August of 2018 and 2019, on Florida Atlantic University’s campus in Jupiter, Florida (26.887596, −80.115436). Caterpillars were collected during the 4^th^ and 5^th^ instar stage. Caterpillars were stored in 16 oz. plastic containers (DELI-36P-16OZ; Comfy Package, Brooklyn, New York), with four caterpillars per container to reflect the density generally observed at the time of collection and generate natural interactions. Caterpillars were fed *A. curassavica* milkweed leaves *ad libitum*; fresh leaves being replaced daily. Plastic containers containing caterpillars were housed in a climate-controlled room at a constant temperature of 25 °C and humidity of 55%, with a 12:12 light/dark cycle. All caterpillars were acclimated to these conditions for at least 1 hour prior to testing. Caterpillars were kept for no longer than three days following collection, and upon completion of behavioral analysis, were released back to the location of their capture.

### Aggression assays

Aggression was measured in arenas containing four caterpillars of the same instar stage. To synchronize feeding, caterpillars were food restricted in an empty container for 1 hr prior to the initiation of each assay. Caterpillars were then transferred to an arena containing fresh milkweed leaves and allowed to acclimate for 1 min prior to testing. Aggressive encounters were recorded using a personal camcorder (Vixia HF R800; Canon, Tokyo, Japan) under standard light conditions over a 10 min period. All data were collected between ZT 5-9:30, where ZT0 represents the time of lights on.

Aggression was determined by manually quantifying the number of aggressive attacks, or lunges, that occurred during each video. Attacks were defined by rapid head movements that make contact with another caterpillar. We partitioned the attack sequence into four actions: (1) recognition, (2) initiation, (3) contact, and (4) cessation. First, the aggressive caterpillar recognizes the recipient caterpillar as a competitor. An aggressive encounter is then initiated upon drawback of the head, followed by a rapid, lateral lunge in the direction of the recipient caterpillar. Next, physical contact between the aggressive caterpillar and recipient caterpillar occurs. Lastly, cessation of an aggressive encounter occurs when the aggressive caterpillar retreats its head from the recipient caterpillar. Incidental contact would occur periodically by two or more caterpillars during food search behavior. An aggressive attack was differentiated by incidental contact by the quick head drawback and lunge movement. The total number of aggressive lunges for each caterpillar was manually counted over the 10-minute period and presented as the mean of all four caterpillars for a single behavioral assay.

### Single animal analysis

Aggressive encounters and related behaviors from individual caterpillars in a representative 10-min video were characterized using the Behavioral Observation Research Interactive Software (BORIS), which allows for single animal analysis through selective coding sequences and focal subject targeting (Friard and Gamba, 2016). Each of four caterpillars were selected as the core subject with the use of BORIS’s parameter controls. With this setting, behavioral coding was used to identify single points (e.g. aggressive encounters) and timed occurrences for each caterpillar (e.g. activity and location metrics).

### Food availability manipulations

We chose amounts of food to reflect variations in food availability that were within the range of naturally occurring conditions, based on our observations of caterpillars at the collection site. At the start of testing, 4^th^ or 5^th^ instar caterpillars were transferred to a petri dish containing low food availability (~ 0.20 g), intermediate food availability (~ 0.30 g) or high food availability (~ 0.42 g) of *A. curassavica*. Caterpillars were allowed to acclimate for a period of 1 minute and then the number of aggressive attacks were quantified, as described above.

### Lighting manipulations

To assess the contribution of light on the number of aggressive encounters, a digital camcorder with a built-in infrared system was used (DCR-SR42; Sony, Tokyo, Japan). These experiments were performed as described above, using 5^th^ instar caterpillars. Caterpillars were first introduced to low food availability with the lights on. Once the lights were turned off, caterpillars were allowed to acclimate for a period of 1 minute, then the number of aggressive attacks were quantified. As a control, similar manipulations were performed, but the lights were never turned off.

### Tentacle ablation

To examine the effect of sensory input on the number of aggressive encounters, the tentacles of 5^th^ instar caterpillars were removed using steel scissors. Tentacles were cut 2–3 mm from the base of each tentacle. Caterpillars were given 24hrs to recover, and then number of aggressive attacks were quantified. Lacerated caterpillars were examined under the same experimental conditions as control caterpillars.

### Statistics

All measurements are presented as bar graphs showing mean ± standard error. A t-test was used for comparisons between two treatments, while a one-way analysis of variance (ANOVA) was used for comparisons between more than two treatments. All post hoc analyses were performed using the Holm-Sidak multiple comparisons test. Statistical analyses and data presentation were performed using InStat software (GraphPad Software 8.0; San Diego, CA).

## Results

To measure aggression in monarch caterpillars, we sought to develop a quantifiable assay for aggression. We first observed caterpillars at the 4^th^ and 5^th^ instar stage, the final stages prior to pupation. In both stages, we noticed a stereotyped aggressive behavior sequence consisting of the aggressive caterpillar orienting towards the conspecific and performing a quick head-snap that made physical contact with the target (Figure 1A,B and Supplemental movie 1). This tended to elicit cessation of feeding and a movement away from jointly occupied space by the target caterpillar. We reasoned that these movements constitute aggression.

**Figure 1.**
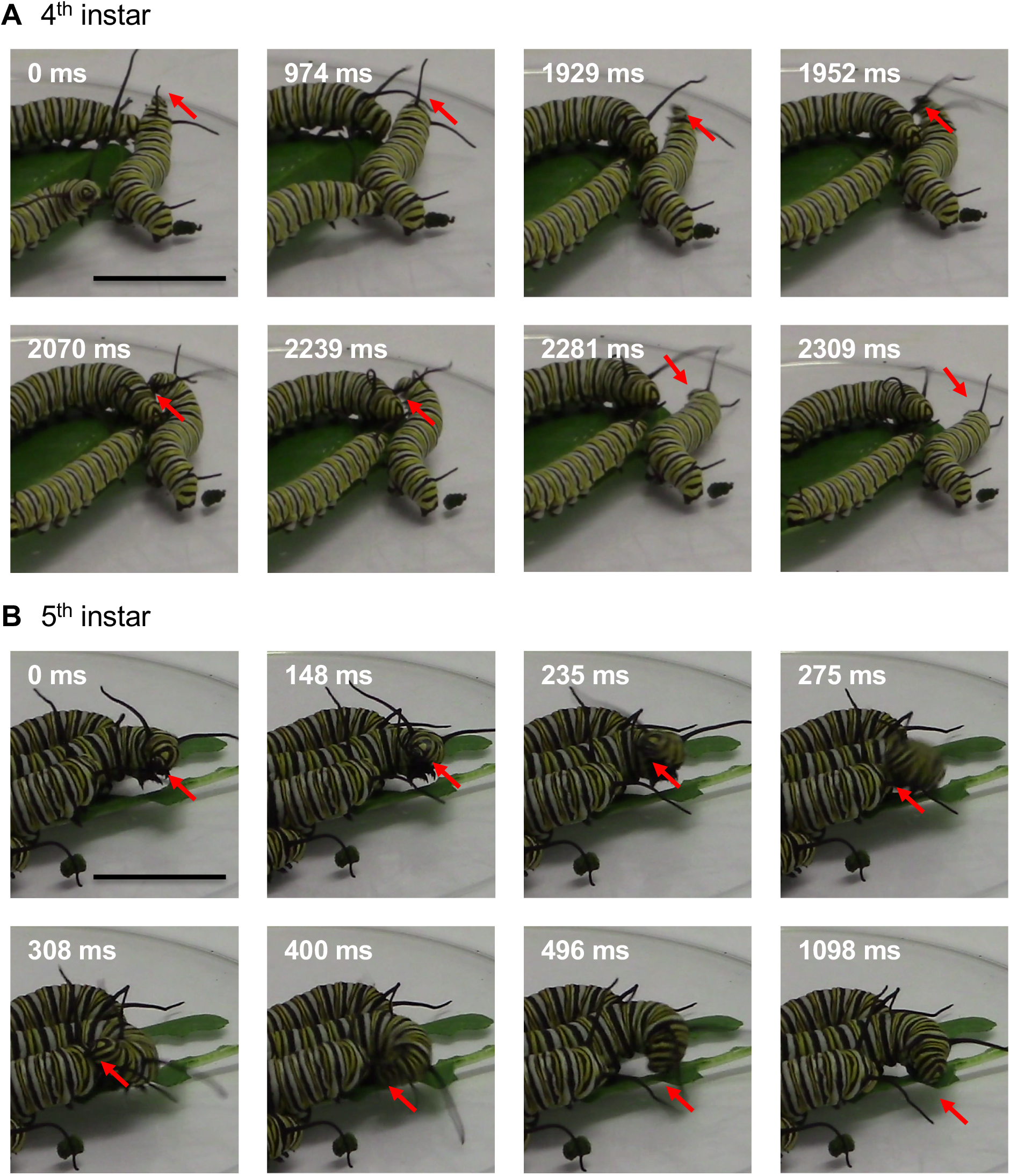
Characterization of aggressive behavior in 4^th^ and 5^th^ instar Monarch caterpillars. Example time-lapse of a stereotypical aggressive encounter, characterized by a lateral lunge, between two (**A**) 4^th^ instar or (**B**) 5^th^ instar conspecifics. Red arrows indicate a change in caterpillar head movement as it progresses from recognition, initiation, contact, and then cessation of a single attack. The timestamp on each image denotes the duration of the encounter, in milliseconds. Scale bar = 2cm.

To determine the relationship between aggression and developmental stage, we quantified the number of attacks across multiple instars. Attacks were not observed in 3^rd^ instar caterpillars (data not shown). A comparison between 4^th^ and 5^th^ instar caterpillars revealed significantly more aggressive lunges between 5^th^ instar caterpillars than between 4^th^ instar caterpillars (Figure 2). Together, these findings suggest aggression increases in frequency over the later stages of larval development. We reasoned that the relatively larger size of older caterpillars may increase competition for food resources, thereby promoting aggression.

**Figure 2.**
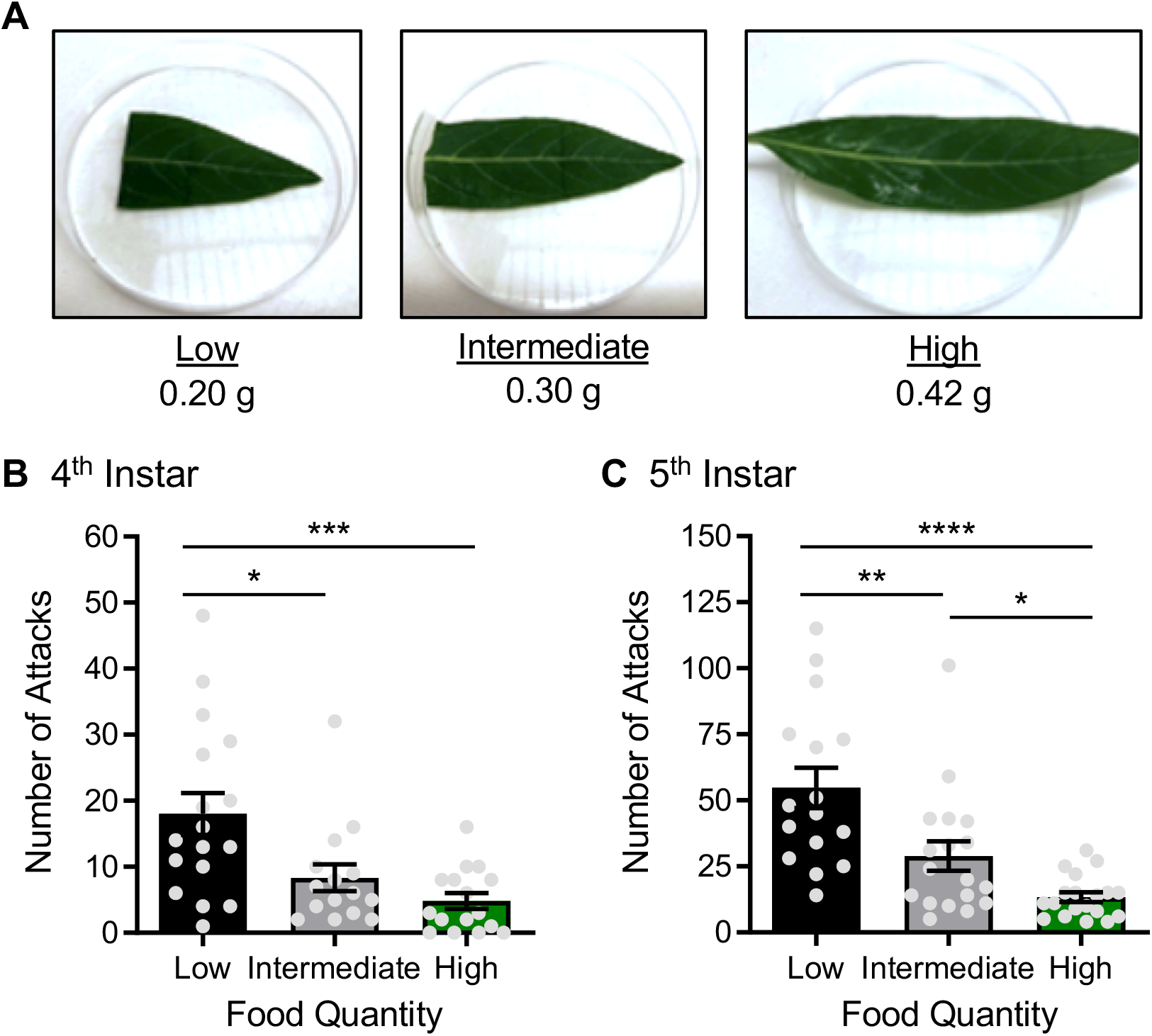
Aggressive behavior increases in both 4^th^ and 5^th^ instar caterpillars with decreasing food density. (**A**) *Asclepias curassavica* of low, intermediate, or high density were provided to 4^th^ and 5^th^ instar caterpillars and aggressive behavior was observed. (**B**) There was a significant effect of food quantity on the number of aggressive attacks in 4^th^ instar caterpillars (F_2, 45_=8.84; *P*<0.0006). *Post hoc* analyses revealed a significant increase in the number of aggressive attacks on low food density relative to both intermediate (*P*<0.0109) and high food densities (*P*<0.0006), while no significant difference between intermediate and high food densities was observed (*P*<0.3002). (C) There was a significant effect of food quantity on the number of aggressive attacks in 5^th^ instar caterpillars (F_2, 50_=15.34; *P*<0.0001). *Post hoc* analyses revealed a significant increase in the number of aggressive attacks on low food density relative to both intermediate (*P*<0.0027) and high food densities (*P*<0.0001). The number of attacks were also significantly higher on intermediate food density in comparison to high food density (*P*<0.0375).

To directly quantify the relationship between food availability and aggression, we generated three conditions that differed in food availability (Figure 2a). Although the total number of attacks were lower among 4^th^ instar caterpillars, there were significantly more attacks among 4^th^ instar caterpillars when food availability was low, with a mean of 18.00 ± 3.17 compared to when food availability was either intermediate or high, with means of 8.33 ± 2.01 and 4.81 ± 1.19, respectively (Figure 2b). The number of attacks among 5^th^ instar caterpillars also correlated with food availability (Figure 2c). We again observed a significantly greater number of aggressive attacks when food availability was low, with a mean of 54.75 ± 7.63, compared to when food availability was either intermediate (28.89 ± 5.55) or high (13.32 ± 1.83). The number of aggressive attacks on intermediate food availability was also greater relative to high food availability. Together, these findings support the notion that low food availability triggers aggression in monarch caterpillars.

To identify when aggression occurs and its impact on the target conspecific, we generated an ethogram of four caterpillars from a single group, defining behaviors (foraging, feeding resting) at the time of each aggressive encounter, and their location (on leaf or off leaf). In nearly all cases, the attacked caterpillar was feeding at the tie of the aggressive encounter, suggesting attacks may disrupt feeding and serve to protect a food source (Figure 3). Similarly, the attacking caterpillar tended to be foraging at the time of the attack, suggesting aggression usually occurs when animals are searching for food, rather than actively consuming food. In addition, we cannot rule out the possibility that crowding plays a role in inducing aggression, as this has been shown to contribute to aggressive behavior in many other insect species (Wang and Anderson, 2010; Stevenson and Rillich, 2016). Therefore, this qualitative analysis supports the notion that aggression is associated with food competition, but other factors are likely involved as well.

**Figure 3.**
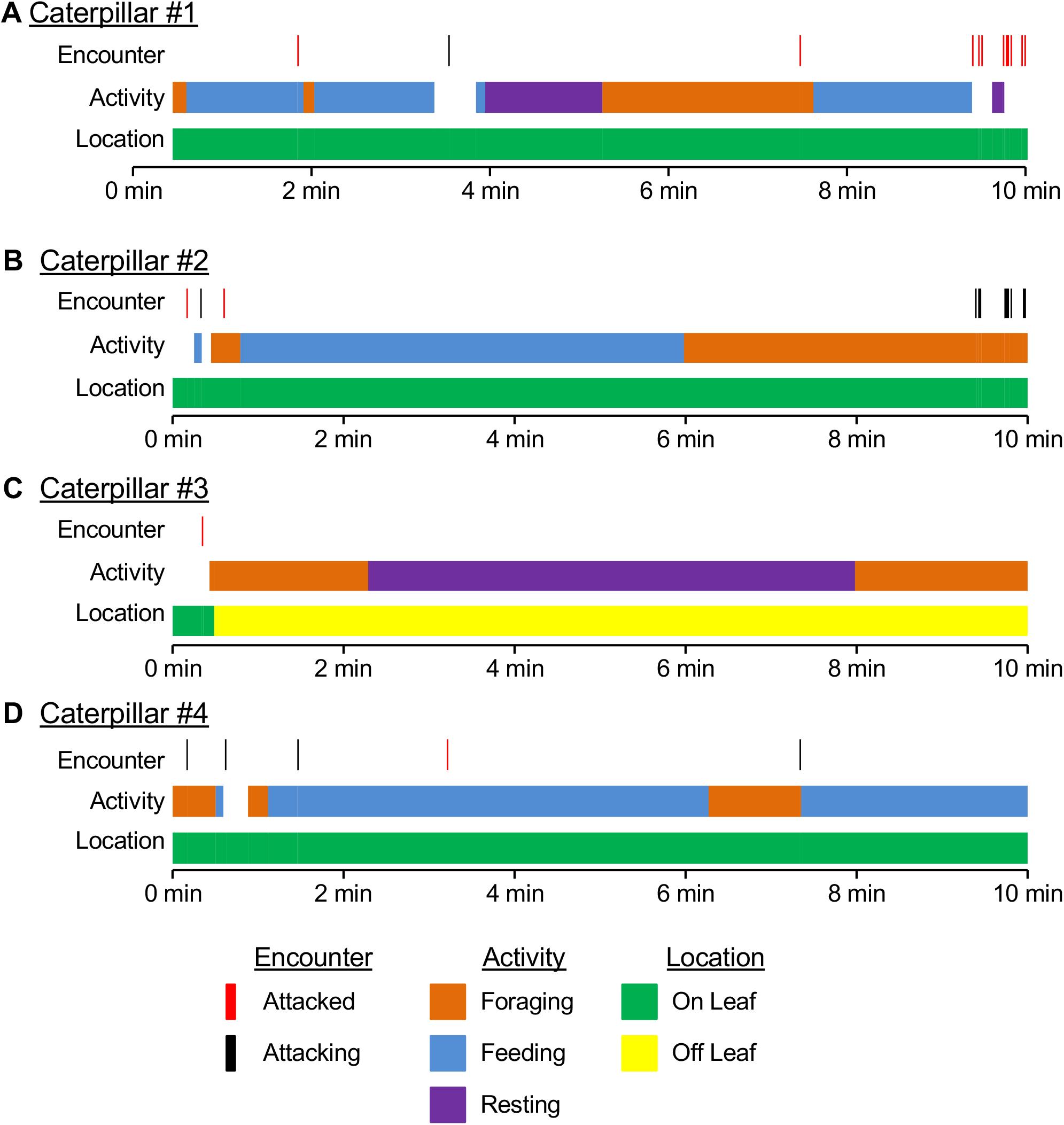
Ethogram of four representative 5^th^ instar caterpillars on intermediate food density during a 10-min video. For each caterpillar, the time at which an aggressive encounter occurs was recorded, as well as its activity and location, shown as fractions of time. Aggressive encounters were classified as either attacking (black) or being attacked (red). Activities were categorized into foraging (orange), feeding (bule), or resting (purple). The location of each caterpillar was also recorded (on the leaf: green; off the leaf: yellow).

To further investigate the defining features of these aggressive attacks, we quantified multiple parameters during each aggressive encounter and its impact on the targeted caterpillar. In both 4^th^ and 5^th^ instars, nearly all aggressive attacks occurred on the leaf (Fig 4a,b). Aggressive attacks were significantly more likely to occur when the target caterpillar was actively feeding and never occurred when a caterpillar was resting (Fig 4c,d). The number of aggressive attacks that occurred during foraging was intermediate between those that occurred during feeding and during resting. Immediately following an attack, the attacking caterpillar almost always remained on the leaf and is consistent for both 4^th^ and 5^th^ instars (Fig 4e,f). In 4^th^ instars, the target caterpillar left the leaf a significantly greater amount of time than it remained (Fig 4e), while results trended in this direction for 5^th^ instar caterpillars (Fig 4f). Therefore, aggression increases the likelihood that the attacking caterpillar will remain on the food source, while the attacked caterpillar will be displaced.

**Figure 4.**
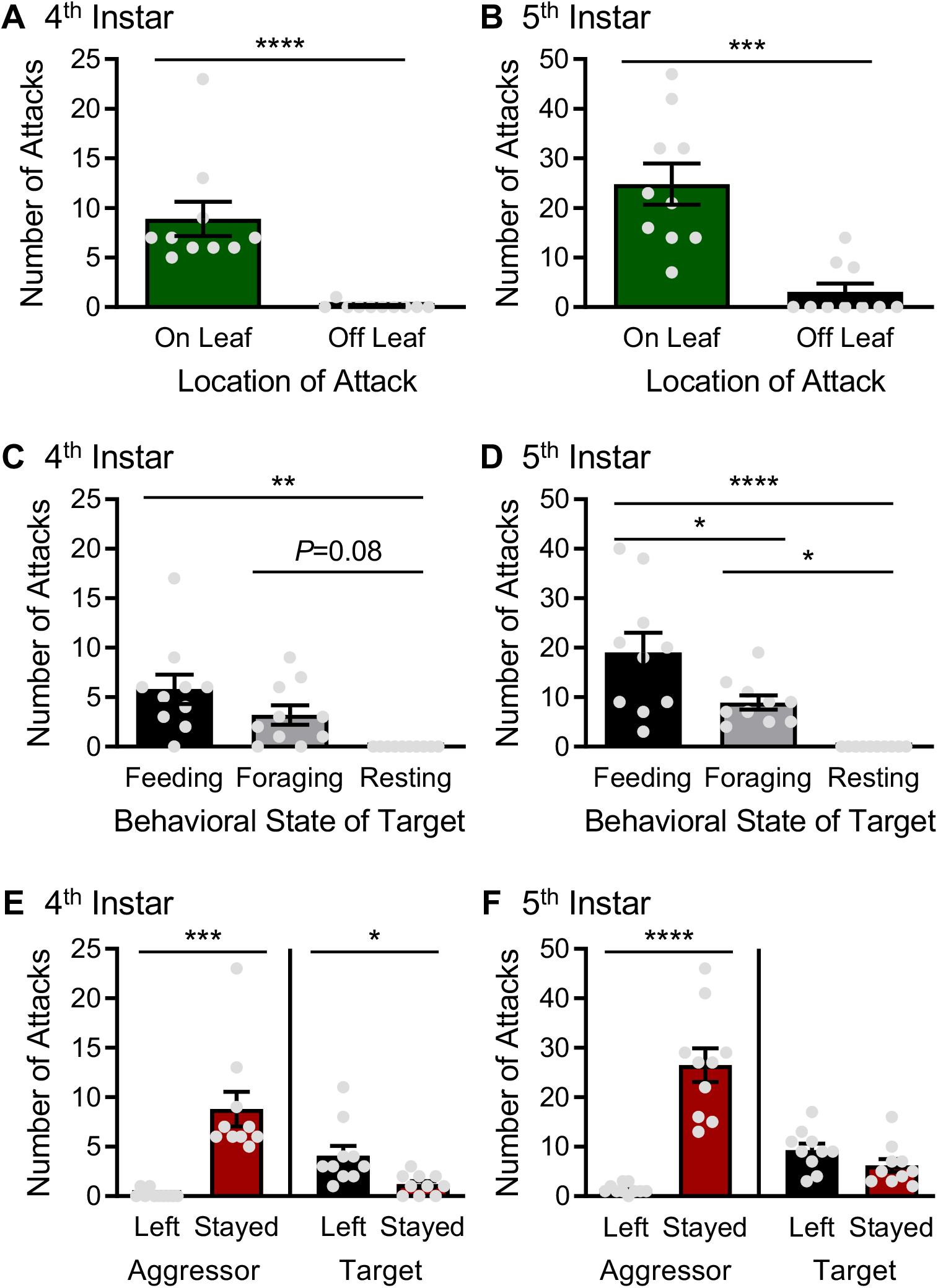
Aggressive behaviors tend to occur on the leaf during feeding. In 4^th^ (**A**) and 5^th^ (**B**) instar caterpillars, there were significantly more aggressive attacks on the leaf than off the leaf (4^th^ instar: F_X, X_=X; *P*<0.X; 5^th^ instar: F_X, X_=X; *P*<0.X). (**C**) There was a significant effect of behavioral state on the number of aggressive attacks in 4^th^ instar caterpillars (F_2,27_=8.047; *P*<0.0018). *Post hoc* analyses revealed that significantly more aggressive attacks occur during feeding relative to resting (*P*<0.0012). No significant difference between feeding and foraging (*P*<0.1903) or between foraging and resting were observed (*P*<0.0876). (**D**) There was a significant effect of behavioral state on the number of aggressive attacks in 5^th^ instar caterpillars (F_2,27_=14.98; *P*<0.0001). *Post hoc* analyses revealed that significantly more aggressive attacks occur during feeding relative to both foraging (*P*<0.0191) and resting (*P*<0.0001). The number of aggressive attacks is also significantly higher during foraging in comparison to during rest (*P*<0.0418). (**G**) At the 4^th^ instar stage, the aggressive caterpillar is significantly more likely to stay on the leaf (t_18_=4.936; *P*<0.0001), while the attacked caterpillar is significantly more likely to leave the leaf (t_18_=2.830; *P*<0.0111). (**H**) At the 5^th^ instar stage, the aggressive caterpillar is significantly more likely to stay on the leaf (t_18_=7.325; *P*<0.0001), but the attacked caterpillar is equally as likely to leave the leaf as they are to stay (t_18_=1.671; *P*<0.1120).

Lastly, we sought to investigate whether sensory input contributes to aggressive behavior. Visually-mediated social behavior has been recently identified in *Drosophila larvae* (Dombrovski *et al.*, 2019), suggesting that simple eyes may convey complex social information. Monarch caterpillars contain six bilateral ocelli allowing for vision (Khan Perveen, 2017). To determine whether visual input is required for aggression, we compared the number of aggressive attacks in 5^th^ instar caterpillars on low food density recorded under standard light conditions, or in complete darkness (Figure 5a). No differences in aggression were identified between the groups (Figure 5b), suggesting light input is dispensable for caterpillar aggression. We also investigated the contributions of the tentacles, large mechanosensory appendages that exhibit rapid growth during the 4^th^ and 5^th^ instar stages when aggression increases (Oberhauser and Kuda, 1997). To investigate the contributions of these structures, we compared aggression in caterpillars with bilateral ablation of the tentacles with intact controls (Figure 5C). There was no difference between the groups, suggesting that functioning tentacles are dispensable for the elicitation of aggression (Fig 4d). These findings suggest that alternative sensory modalities, such as pheromonal, olfactory, or tactile cues that are independent of the tentacles initiate aggression.

**Figure 5.**
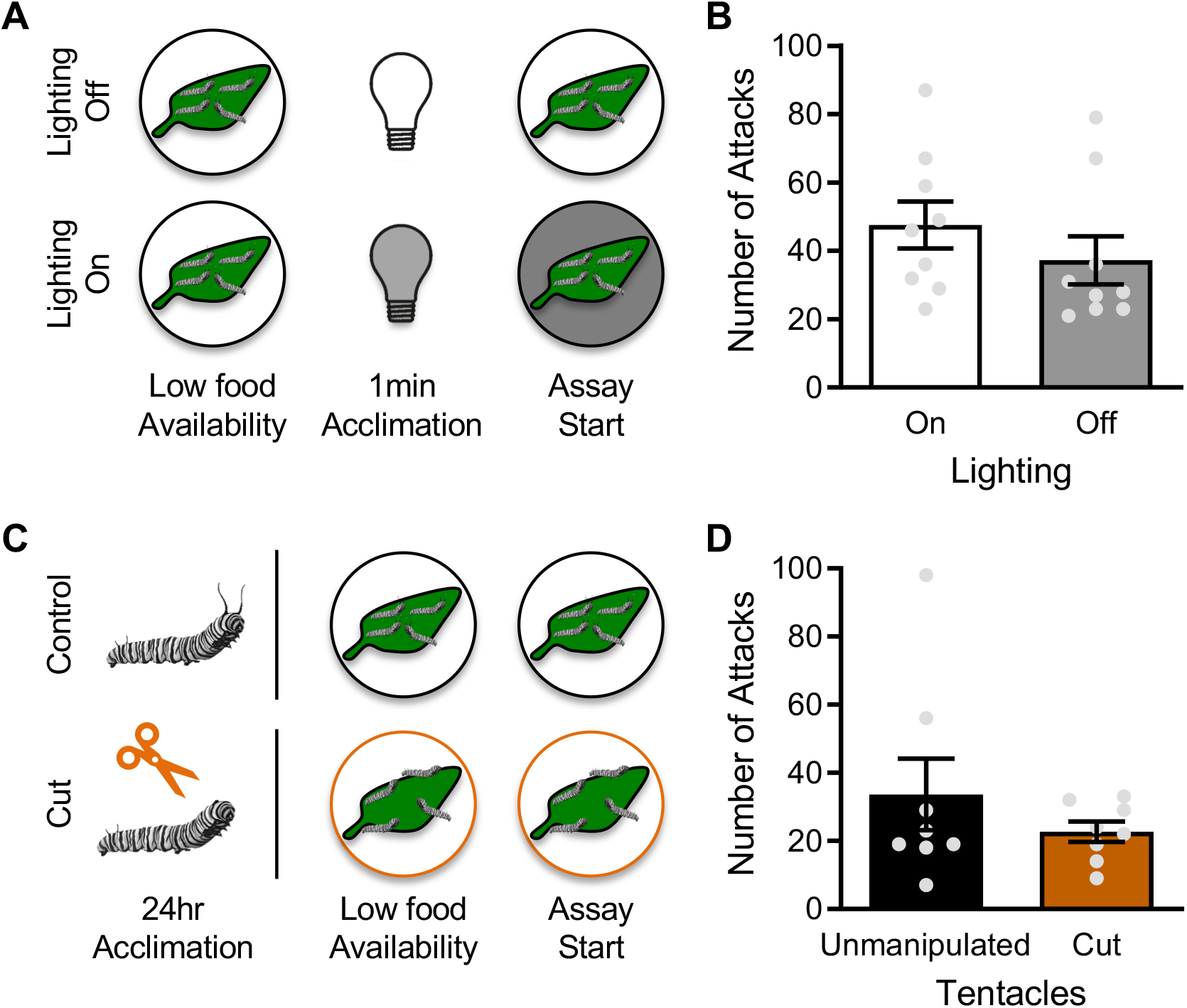
Lighting and tentacle laceration have no effect on aggressive behavior. (A) In the experimental treatment, lights were turned off 1 min prior to testing. The effect of lighting on aggressive behavior was then measured in arenas with low food availability. (B) Caterpillars were equally as aggressive when the lights are off as they are in lighted conditions (t_16_ = 1.054. *P*<0.3076). (C) In the experimental treatment, tentacles from each caterpillar were removed 24 hrs prior to testing. The effect of tentacle laceration was then measured in arenas with low food availability. (D) Caterpillars were equally as aggressive without tentacles as they are with intact tentacles (t_14_ = 0.9964. *P*<0.3360).

## Discussion

Here, we describe aggressive behavior in a caterpillar that consists of physical contact presumably to defend a food source that is critical for development. The physical contact is more likely to occur on food and frequently results in displacement suggesting it may be related to resource acquisition. It is important to note that we observed a high degree of variability between individuals, raising the possibility that interindividual differences in exploration, social behavior, or boldness underlie response differences. Territoriality has been described in other caterpillar species (Kemp, 2000; Yack, Smith and Weatherhead, 2001; Bowen *et al.*, 2008), suggesting aggression may be widespread. Cannibalism is widespread in *Lepidoptera* larvae (Semlitsch and West, 1988; Dial and Adler, 1990; Zago-Braga and Zucoloto, 2004; Tang *et al.*, 2016; Zhou, Dudash and Fenster, 2016), raising the possibility that aggression is associated with conspecific competition. This extends beyond *Lepidoptera* to other insect larvae, as crowded rearing conditions promote cannibalism in *Drosophila* larvae (Vijendravarma, Narasimha and Kawecki, 2013), suggesting a relationship between resource availability and this aggression-linked behavior. In our study, we exclusively examined behavioral lunges, a feature most commonly associated with aggressive behavior (Dankert *et al.*, 2009). Examining acoustic communication and potential cannibalism in monarch caterpillars may inform additional behavioral components associated with resource competition.

Numerous sources of resource competition trigger aggression, including limited availability of territory, mates, and food. Our findings suggest limited food availability triggers aggression in monarch caterpillars. In contrast, in social species like the honeybee, *Apis mellifera*, rather than an increase in aggression, nutritional stress during larval development promotes cooperation, in which the queen increases colony investment and decreases reproduction (Walton *et al.*, 2018). However, the effect of nutritional state on aggression is likely wide-spread throughout the animal kingdom as food scarcity induces aggression in amphibians, lizards, and birds (Moore and Marler, 1987; Drummond and Garcia Chavelas, 1989; Ducey and Heuer, 1991). In facultatively sublicidal bird species, where aggression commonly results in death of the competitor, the “food amount hypothesis” contends that sibling aggression will vary inversely with food quantity (Mock, Lamey and Ploger, 1987; Drummond, 2001a). Although less is known about the proximate causes of aggression in nonsiblicidal species (Drummond, 2001b), in the meerkat, *Suricata suricatta*, the severity of aggressive competition scales with food availability (Hodge *et al.*, 2009). Similarly, in numerous mammalian models, moderate reductions in food availability promote aggression. For example in Siberian hamsters, *Phodopus sungorus*, food restriction promotes aggression during winter-like photoperiods, a signal of impending limited resources (Hodge *et al.*, 2009; Allison M. Bailey, Nikki M. Rendon, Kyle J. O’Malley, 2017). Food restriction in utero is also associated with aggression. In baboons, *Papio hamadryas*, interauterine growth restriction is linked to higher rates of aggressive display behaviors such as aggressive grunts, canine displays, and eyelid flashes (Huber *et al.*, 2015). In *Drosophila*, which is widely used to study aggression in the laboratory, conspecific aggression is triggered by placing a small amount of high calorie food in the center of an arena (Chen *et al.*, 2002; Lim *et al.*, 2014). Therefore, we hypothesize that aggression induced by limited food availability in monarch caterpillars are likely present in many different species throughout the animal kingdom.

While we systematically tested the relationship between food availability and aggression, we cannot exclude other possibilities. For example, it is possible that low amounts of food brings caterpillars into closer proximity, thereby resulting in aggression. In this case, it is proximity, rather than food availability *per se* that induces aggression. The conditions, all of which included different amounts of food were chosen to mimic conditions in the wild. The complete lack of food resulted in roaming behavior, where caterpillars are highly active. This likely represents a foraging state, and few aggressive events were observed (data not shown). Manipulating feeding state prior to testing may provide additional insight into the relationship between feeding state and aggression.

In most species studied, aggression is dependent on multiple sensory inputs, providing the opportunity to identify specified sensory stimuli that induce aggression (Chen and Hong, 2018). For example in flies, dedicated pheromone sensing neurons, and their cognate receptors, are required for aggression (Wang and Anderson, 2010). Further, disruption of the visual system also disrupts aggressive behavior (Hoyer *et al.*, 2008; Ramin *et al.*, 2014). Conversely, we find that aggression is similar monarch’s under light and dark conditions, suggesting it occurs independently of visual function. Further, ablation of the tentacles does not impact aggression, suggesting mechanosensory stimuli from these organs are not required. It is possible that other sensory modalities, such has acoustic communication contribute to the regulation of aggression. For example, in the warty birch caterpillar, *Drepana bilineata*, acoustic cues are used to defend silk leaf mats and shelters from conspecific larvae (Bowen *et al.*, 2008). Further, caterpillars of the silk moth family *Bombycoidea* display a wide array of defense sounds in response to simulated inter-species aggression (Bura, Kawahara and Yack, 2016). Future work examining the contributions of olfactory or alternative mechanosensory pathways may help identify neural mechanisms that trigger aggression.

Monarch caterpillars are specialists that feed nearly exclusively on the milkweed (Seiber *et al.*, 1986; Malcolm, Cockrell and Brower, 1989). It is likely that food resources are scarcer for specialists than generalists, or during life stages where animals are dietary specialists, raising the possibility that conspecific aggression is elevated in this species. Food competition may be more prevalent for monarch caterpillars that largely defoliate host plants, generating resource scarcity, particularly in the later stages of larval growth when we observed aggression (Kaplan and Denno, 2007). In addition, it is possible that conspecific aggression originated as a mechanism of inter-species defense. Over 100 different species are known to consume neonate monarch caterpillars (Hermann *et al.*, 2019). Investigating the conditions that trigger aggression in the wild, and in additional *Lepidotera* caterpillars, will allow for comparative analysis that may provide insight into the ecological factors that promote aggression.

Beyond the study of aggression in caterpillars, monarchs present an emerging model for studying the molecular mechanisms underlying behavior. Monarch caterpillars are widely studied for their migratory behavior that spans generations along the animals range from Southern Mexico through Canada (Brower, 1996; Reppert, Gegear and Merlin, 2010). Monarch butterflies detect polarized light and use a time-compensated sun compass to for migratory navigation (Froy *et al.*, 2003; Reppert, Zhu and White, 2004). The genome of monarch butterflies has been sequenced and the recent implementation of gene editing provides the opportunity to investigate the genetic mechanisms underlying behavior (Zhan *et al.*, 2011; Merlin *et al.*, 2013; Markert *et al.*, 2016). Our findings that monarch caterpillars exhibited aggressive behavior under conditions of limited food availability provide the opportunity to examine the role of genes identified in other leading models of aggression in an ecologically relevant system.

## Acknowledgements

This work was supported by NSF awards DEB 174231 and IOS 165674 to ACK. We are grateful for technical support from Christine Merlin (Texas A&M) and for guidance on establishing aggression assays from Kenta Asahina (Salk Institute).

